# NONLINEAR NEURAL NETWORK DYNAMICS ACCOUNTS FOR HUMAN CONFIDENCE IN A SEQUENCE OF PERCEPTUAL DECISIONS

**DOI:** 10.1101/648022

**Authors:** Kevin Berlemont, Jean-Rémy Martin, Jérôme Sackur, Jean-Pierre Nadal

## Abstract

Electrophysiological recordings during perceptual decision tasks in monkeys suggest that the degree of confidence in a decision is based on a simple neural signal produced by the neural decision process. Attractor neural networks provide an appropriate biophysical modeling framework, and account for the experimental results very well. However, it remains unclear whether attractor neural networks can account for confidence reports in humans. We present the results from an experiment in which participants are asked to perform an orientation discrimination task, followed by a confidence judgment. Here we show that an attractor neural network model quantitatively reproduces, for each participant, the relations between accuracy, response times and confidence. We show that the attractor neural network also accounts for confidence-specific sequential effects observed in the experiment (participants are faster on trials following high confidence trials). Remarkably, this is obtained as an inevitable outcome of the network dynamics, without any feedback specific to the previous decision (that would result in, e.g., a change in the model parameters before the onset of the next trial). Our results thus suggest that a metacognitive process such as confidence in one’s decision is linked to the intrinsically nonlinear dynamics of the decision-making neural network.

## Introduction

A general understanding of the notion of confidence is that it quantifies the degree of belief in a decision Meyniel et al. [2015b], Mamassian [2015]. The simplest context for studying confidence is the one of perceptual decision making. In psychology and neuroscience, the most commonly used experimental protocols are the ones of two alternative forced choices (2AFC) and stimulus discrimination tasks in which, in a sequence of trials, the participant is presented with stimuli and has to make a binary choice, associating each stimulus to one of two categories (e. g. decide if a visual stimulus is the image of a cat or a dog). Many studies have tackled the issue of confidence measurement in perceptual decision tasks, either by directly requiring participants to provide an estimation of their confidence Peirce and Jastrow [1884], Zylberberg et al. [2012], Adler and Ma [2018], or by using postdecision wagering (subjects can choose a safe option, with low reward regardless of the correct choice) Vickers [1979 (reeditited in 2014], Fleming et al. [2010], Kepecs and Mainen [2012]. Postdecision wagering has been used in behaving animals in order to study the neural basis of confidence Kepecs et al. [2008], Kiani and Shadlen [2009], Komura et al. [2013], Lak et al. [2014].

In order to model the neural mechanisms underlying the decision-making process, two main routes are followed. The most frequently used considers (linear) drift-diffusion models (DDM) Bogacz et al. [2006], Ratcliff [1978] or independent race models (IRM) Raab [1962], Vickers [1970], Merkle and Van Zandt [2006], in which choice-specific cells accumulate evidence in favor of one or the other alternative to which they are tuned. A more biophysical approach considers attractor neural networks Wang [2002], with competing pools of cells, leading to a nonlinear dynamics with choice-specific attractors. Within one or the other framework, researchers have tried to relate confidence to the decision-making process, making different hypotheses on the origin of confidence. Within the Bayesian and signal detection theories, researchers model confidence from the probability of having made the correct choice Clarke et al. [1959], Fleming et al. [2010], Kepecs and Mainen [2012], Yeung and Summerfield [2012], Meyniel et al. [2015a]. When considering the neural dynamics, researchers assume that confidence is based on the integration of evidence over time Kepecs et al. [2008], Pleskac and Busemeyer [2010], Wei and Wang [2015]. Finally, researchers have modeled confidence as based on a consensus reached by a pool of independent decision-making networksKoriat [2012], Paz et al. [2016].

Several experimental studies Kepecs and Mainen [2012], Meyniel et al. [2015b] suggest that choice and confidence can be read out from the same neural representation. In an experiment by Kiani and Shadlen Kiani and Shadlen [2009], monkeys perform a discrimination task (each correct choice leading to a reward), but on half of the trials, the monkey is given the option to abort the task in favor of a certain but small reward. The probability of choosing this ’sure target’ reflects the monkey’s degree of choice uncertainty, assuming that risk aversion strongly correlates with this uncertainty. In order to account for the experimental findings in this uncertain option task, authors Wei and Wang [2015], Jaramillo et al. [2019] have proposed biophysical attractor neural network models. They show that these models capture both the behavioral observations and the associated physiological recordings from neurons in the Lateral Intraparietal (LIP) cortex, area where the firing rates of individual neurons strongly correlate with the decision that is being made, and in the pulvinar area, which might be the locus of the readout of confidence from the LIP activity.

In the present work, we address the issue of the ability of attractor networks to quantitatively account for confidence reports in human. For this, we first experimentally investigate confidence formation and its impact on sequential effects in human experiments. Participants perform an orientation discrimination task on Gabor patches that deviate clockwise or counter-clockwise with respect to the vertical. In some blocks, after reporting their decisions, participants perform a confidence judgment on a visual scale. Then, we fit an attractor neural network model Wong and Wang [2006], Berlemont and Nadal [2019] on the behavioral data. More precisely, for each participant, we calibrate a network specifically on his/her behavioral data, the fit being only based on mean response times and accuracy. With the model so calibrated for each participant, and making simulations that replicate the experimental protocol, here for the first time we quantitatively confront an attractor neural network behavior with human behavior during full sequences of perceptual decisions. Following Wei and WangWei and Wang [2015], we assume that confidence is an increasing function of the difference, measured at the time the decision is made, between the mean spike rates of the two neural pools specific to one or the other of the two possible choices. We show that in this way, behavioral effects of confidence can be accurately estimated for each participant. We find that the attractor neural network accurately reproduces an effect of confidence on serial dependence which is observed in the experiment: participants are faster (respectively slower) on trials following high (resp. low) confidence trials. Since drift diffusion models cannot account for such effects without ad-hoc changes of parameters from trial to trial, we argue that these sequential effects reveal the intrinsically non-linear nature of the underlying neural network dynamics.

## Results

### Experiment and neural model

Participants completed a visual discrimination task between clockwise and counter-clockwise orientated stimuli, followed, or not, by a task in which they were asked to assess the confidence in their decision. We used three kinds of blocks, comprising either sequences of pure decision trials (*pure* blocks), trials with feedback (*feedback* blocks) or trials with confidence judgments (*confidence* blocks). In *feedback* blocks, on each trial, participants received auditory feedback on the correctness of their choice. In the *confidence* block, after each trial, they were asked to report their confidence on a discrete scale of ten levels, from 0 to 9. In *feedback* blocks, participants were not asked to report their confidence, and in *confidence* blocks they did not receive any feedback. We illustrate the experimental protocol in Fig. 1, panels A - D.

**Figure 1:**
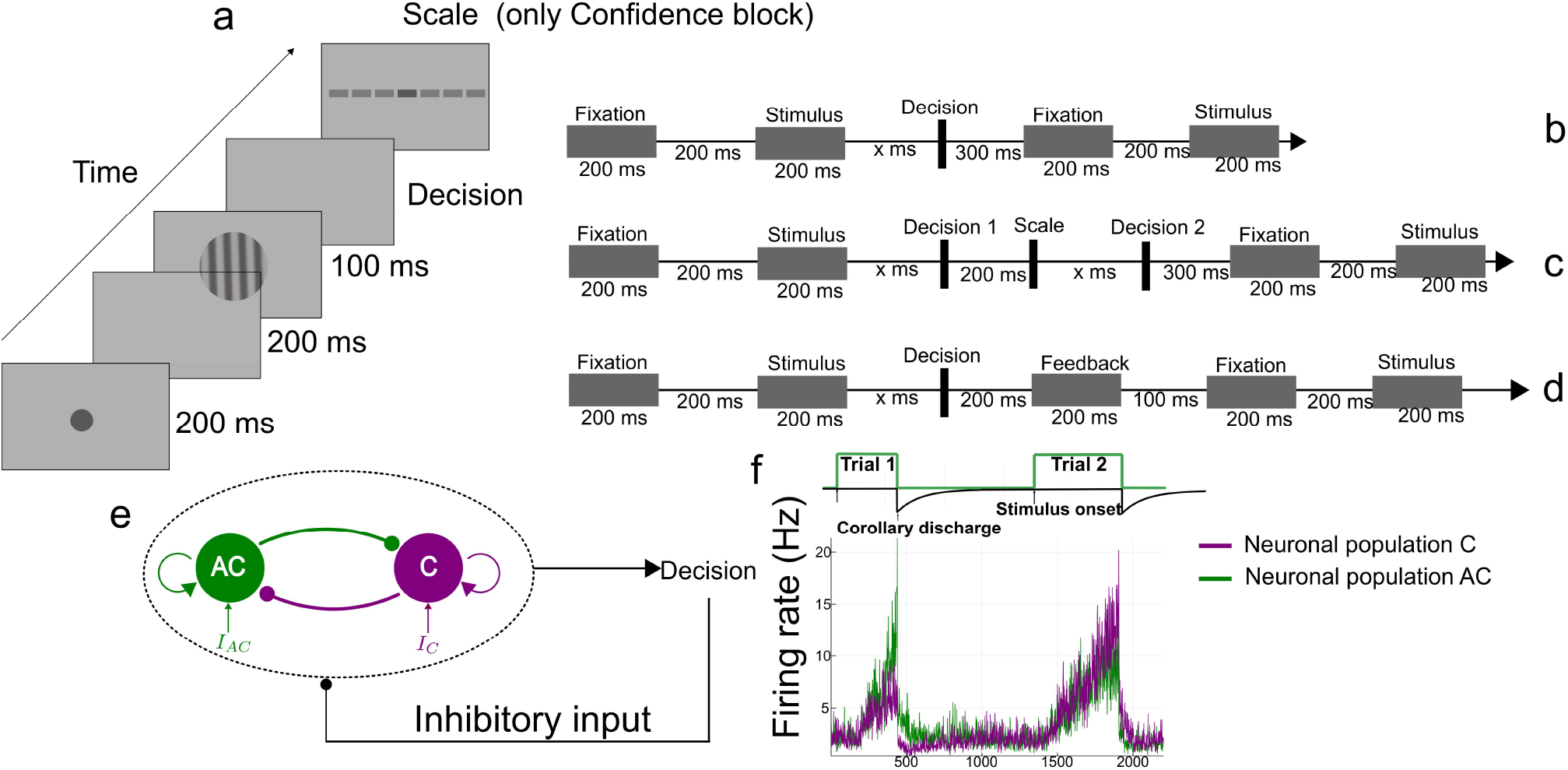
Experimental protocol and model architecture. Procedure of the discrimination task, for the three blocks. (A) Structure of a trial: Following a fixation period, the circular grating (Gabor patch, oriented clockwise, C, or counterclockwise, AC) appears and participants make the decision (C or AC). In confidence blocks, after a delay, participants report their confidence with respect to their choice, on a discrete scale with 10 levels. (B) Time course of a pure block trial. (C) Time course of a confidence block trial. (D) Time course of a feedback block trial. (E) Decision-making network structure. The network consists of two neural pools (specific to clockwise (C) and anti-clockwise (AC) stimuli), endowed with self-excitation and mutual inhibition. After a decision is made (threshold crossed), a non specific inhibitory input (corollary discharge) is sent onto both units. (F) Time course of the neural activities of both pools during two consecutive trials.

For the modelling of the neural correlates, we consider a decision-making recurrent neural network governed by local excitation and feedback inhibition, based on a biophysical model of spiking neurons Compte et al. [2000], Wang [2002]. We work with a reduced version Wong and Wang [2006] allowing for large-scale numerical simulations and for better analytic analysis. More precisely, we consider a model variant Berlemont and Nadal [2019] allowing the network to engage in a sequence of perceptual decisions, as explained below.

The model (see Fig. 1.E) consists of two competing units, each one representing an excitatory neuronal pool, selective to one of the two available response options, here *C* (clockwise) or *AC* (anti-clockwise). Each population receives a task-related input signaling the perceived evidence for each option. The difference between these inputs varies inversely with the difficulty of the task, thus it varies with the absolute value of the Gabor orientation. The decision, ’*C*’ or ’*AC*’, is made when one of the two units reaches a threshold *z*. Once a decision is made (threshold is reached), a non specific inhibitory current (the corollary discharge) is injected into the two neural pools, causing a relaxation of the network activity towards a neutral low activity state, before the onset of the next stimulus. This allows the network to deal with consecutive sequences of trials, as illustrated in Fig. 1.F. For a biologically relevant range of parameters, relaxation is not complete at the onset of the next stimulus, hence the decision made in this new trial will depend on the one at the previous trial. In a previous work, we showed Berlemont and Nadal [2019] that the model accounts for the main sequential and post-error effects observed in perceptual decision making experiments in human and monkeys.

Full details about the experiment and the model can be found in the Methods Section.

### Calibration of the model onto the behavioral results

In Fig. 2, we show for each participant response times and accuracies with respect to stimulus orientation (absolute value of the orientation angle). All subjects exhibit improved accuracies and shorter response times for less difficult (larger orientation) stimuli, as classically reported in the literature.

**Figure 2:**
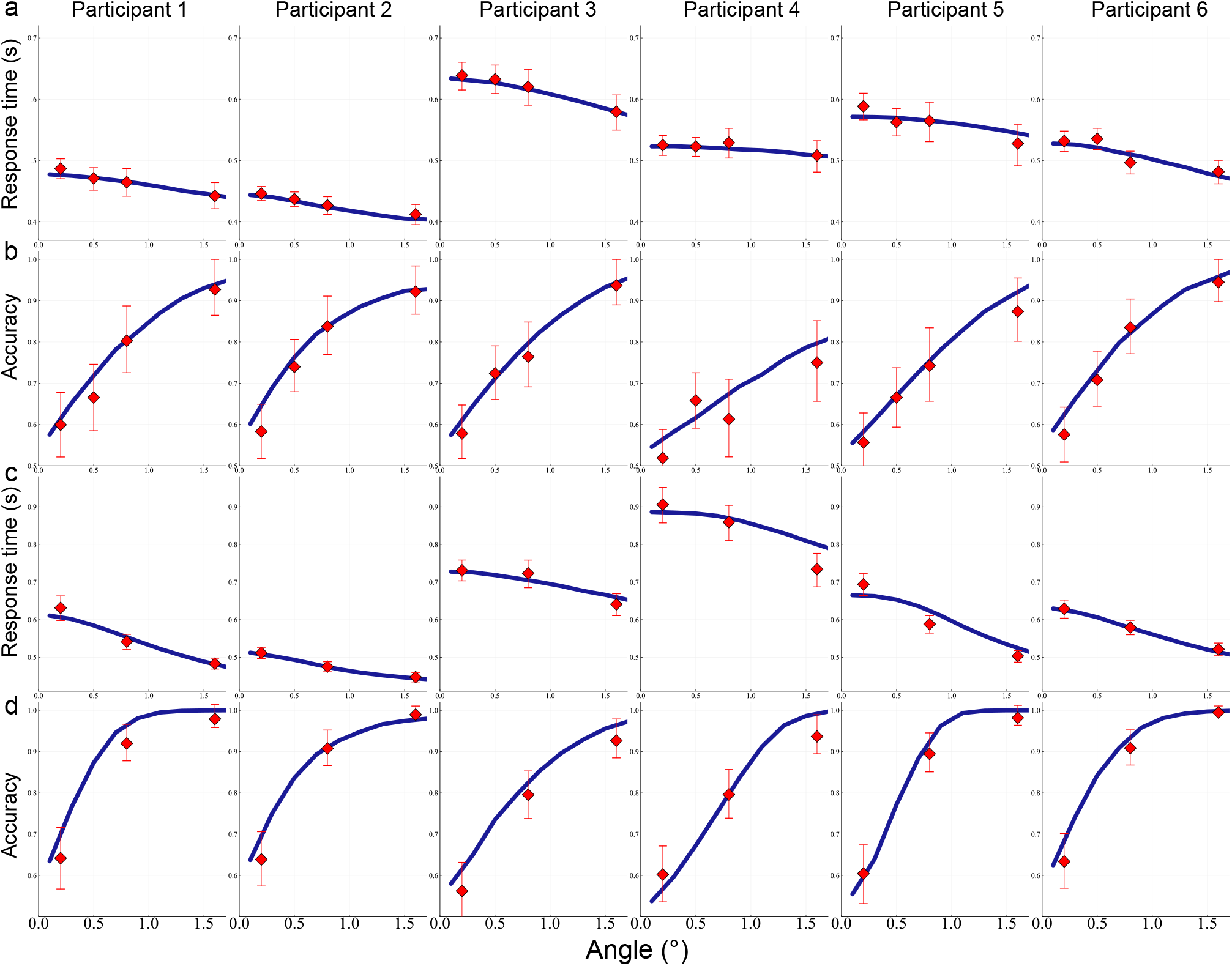
Mean response times (A,C) and accuracies (B,D) as a function of the absolute value of stimulus orientation, *in the* pure *(A and B) and* confidence *(C and D) blocks. For each subject we represent the behavioral data (red dots) and the associated fitted model (blue line). Error bars are* 95% *confidence interval using the bootstrap method*.

We fit the model to these behavioral data. As detailed in the Methods Section, we perform model calibration in order to reproduce both the mean response times and the accuracy (success rates). For each participant, this is done separately for the three types of blocks.

First, we note that the model correctly reproduces the behavioral results of the different participants, as can be seen in Fig. 2. Second, we compare the values of the parameters obtained for the pure and confidence blocks. We find that participants have higher decision threshold (Signed Rank test Wilcoxon [1945] *p* = 0.03, 6 participants), higher stimulus strength level by angle (Signed Rank test, *p* = 0.031, 6 participants) and higher mean non-decision times (Signed Rank test *p* = 0.03, 6 participants). Two of the authors of the present paper (*J-R. Martin, J. Sackur, personnal communication, April, 2018*) have obtained analogous results when analyzing similar data within the Drift Diffusion framework: nondecision time, drift rate and decision threshold are modified by the confidence context in the experimental setup.

### Non-decision times

Our fitting procedure allows estimating the non-decision times. In Fig. 3, we represent the histogram of the response times across participants for the pure and confidence blocks. The red curve shows the distribution of non-decision times in the model, and the black curve the response times distribution. We note that, with a fit only based on the *mean* response times and accuracies, the model also accurately account for the *distributions* of response times. We find that the minimum value of non-decision time is 75 ms for the pure block, and 100 ms for the confidence block, and the average non-decision times are within the order of magnitude of saccadic latency Luce et al. [1986]. Finally, we observe that the non-decision times distributions clearly show a right skew for several participants, in agreement with previous studies Verdonck and Tuerlinckx [2016]. This justifies the modelling of non-decision times with an exponentially modified Gaussian (ex-Gaussian) distribution, instead of simply adding a constant non-decision time to every decision time.

**Figure 3:**
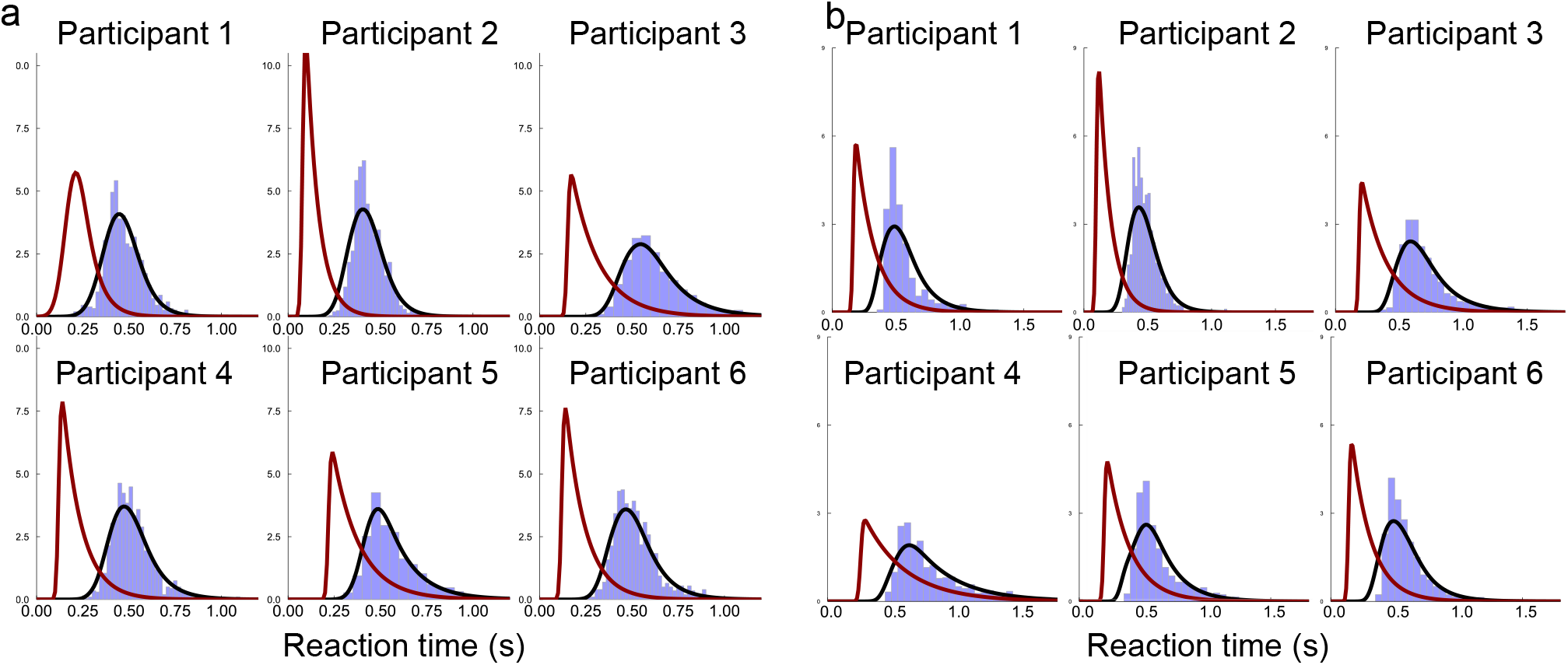
Distributions of RTs for each subject. *(A)* Pure *block data, (B)* confidence block *data. For both panels: In blue, participants’ histograms of the response times; Black curve: density of response times of the simulated network model; Red curve: the associated non decision response times distribution*.

## Confidence modeling

Recent studies have reported a choice-independent representation of confidence signal in monkeys Ding and Gold [2011] and in rats Kepecs et al. [2008], as well as evidence for a close link between decision variable and confidence – in monkeys from LIP recordings Kiani and Shadlen [2009] and in humans from fMRI experiments Hebart et al. [2014]. In an experiment with monkeys,Shadlen Kiani and Shadlen [2009] introduce a ’sure target’ associated with a low reward, which can be chosen instead of the categorical targets. The probability of *not* choosing the sure target is then a proxy for the confidence level. Wei and Wang Wei and Wang [2015] model the neural correlates of confidence within the framework of attractor neural networks. They assume that the confidence level (as given by the probability of not choosing the sure target) is a sigmoidal function of the difference, at the time of decision, between the activities of the winning and loosing neural pools. This hypothesis is in line with similar hypothesis in the framework of DDMs and other decision-making models Vickers [1979 (reeditited in 2014], Mamassian [2015]. They then show that the empirical dependencies of response times and accuracies in the confidence level are qualitatively reproduced in the simulations of the neural model.

We make here the hypothesis that the confidence in a decision is based on the difference ∆*r* between the neural activities of the winning and loosing neural pools Wei and Wang [2015], measured at the time of the decision: the larger the difference, the greater the confidence. In our experiment, the measure of confidence is the one reported by the subjects on a discrete scale, and it is this reported confidence level that we want to model. Within our framework, we quantitatively link this empirical confidence to the neural difference ∆*r* by matching the distribution of the neural evidence balance with the empirical histogram of the confidence levels. In Fig. 4, we show, for each participant, the matching between the histogram of confidence levels, as reported by the participant, and the distribution of ∆*r*, as obtained in the model calibrated on the participant performance. We note that the main difference between the participants’ histograms lies in the percent of trials (and level on the confidence scale) for which a participant reports the highest confidence level. This last point is highly dependent on the participant, and can be at a very low value of ∆*r* (see e.g. Participant 2 on Fig. 4.B).

**Figure 4:**
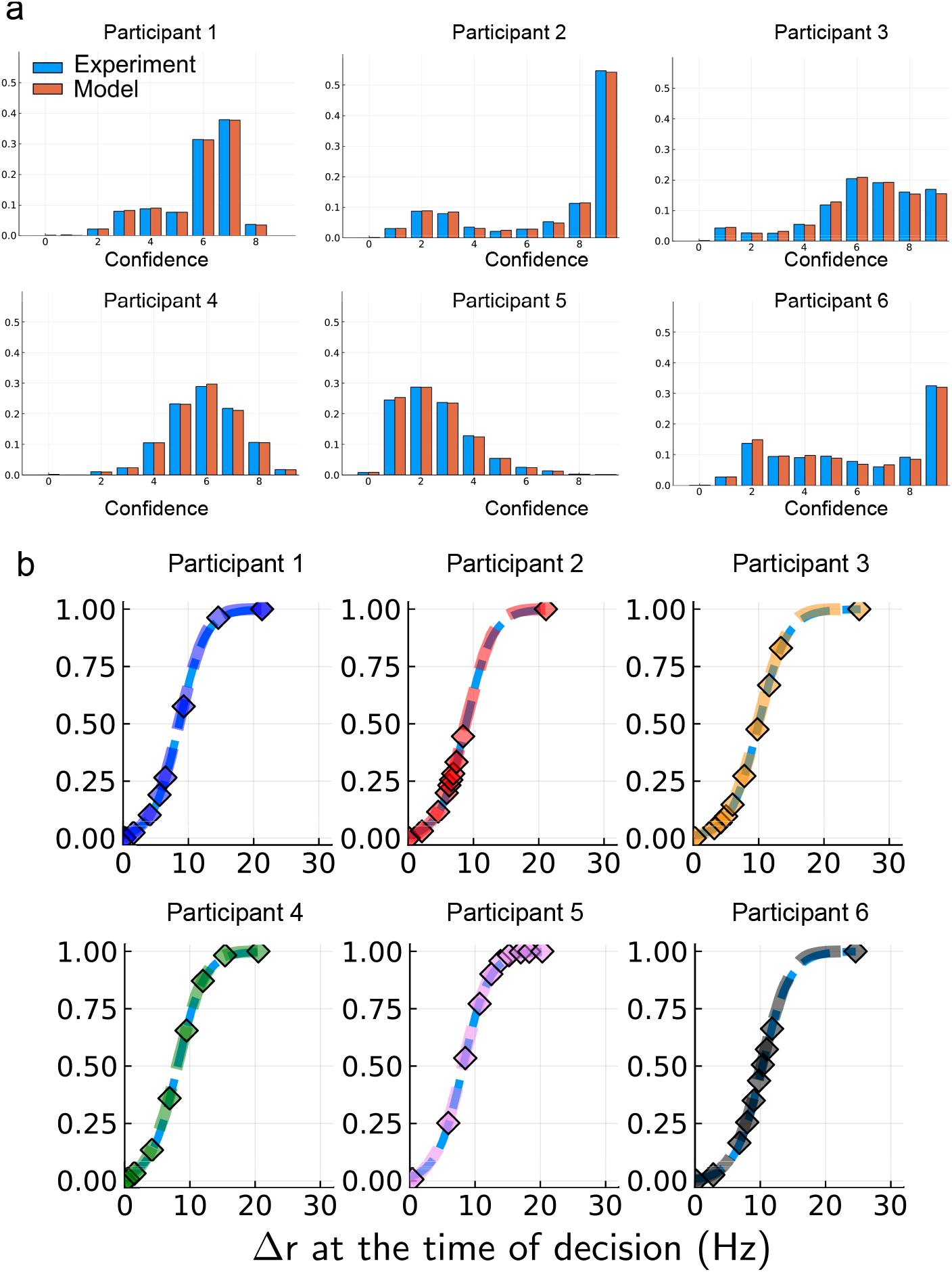
Matching network confidence measure to empirical behavioral confidence. *(A) Confidence histograms. The x-axis gives the value of the confidence on a discrete scale from 0 to 9. Each sub-panel corresponds to a different participant with, in blue, the histogram of the reported confidence, and in orange, the one from the model. For clarity we plot the blue and orange bars side by side, but the bins of the histograms are, by construction, identical. (B) Transfer function F for each participant. The x-axis denotes the difference in neural pools activities* ∆*r at the time of the decision, and the y-axis the cumulative distribution of* ∆*r. Each point represents the levels of* ∆*r* delimiting the level of confidence (from left to right, confidence level 0 to confidence level 9). The dashed colored curve is the cumulative distribution function (CDF) and the light blue dashed curve is the fit of the CDF by a sigmoid.

In our analysis, the shape of the mapping is not chosen a priori but non-parametrically inferred from the experimental data. This is in contrast with previous studies in which the sigmoidal shape is imposed Beck et al. [2008], Kepecs et al. [2008], Kepecs and Mainen [2012]. In a related neural attractor model, Wei and WangWei and Wang [2015] exhibit a link between ∆*r* and a probabilistic measure of confidence, and show that it is well fitted by a sigmoid function. Here we find that, for each participant, the non-parametric mapping is also very well approximated by a sigmoidal function of the type 1*/* (1 + exp (*β*(∆*r − κ*))), with participant-specific parameters *κ* and *β*. It is noticeable that both the link between ∆*r* and a probabilistic measure of confidence observed in an attractor network model, and the mapping obtained here between ∆*r* and the empirical confidence, can be approximated by a sigmoid function. This suggests that the human reported confidence can be understood as a discretization of a probabilistic function.

### Response times and accuracies vs. Confidence

Studies have shown that confidence ratings are closely linked to response times Baranski and Petrusic [1994], Desender et al. [2018a] and choice accuracy Baranski and Petrusic [1994], Sanders et al. [2016], Urai et al. [2017]. The behavioral confidence in our model is assumed to be based on a simple neural quantity measured at the time of the decision. In what follows, we study whether this hypothesis on the neural correlates of confidence can account for the links between the behavioral data: response times, accuracy and confidence. In Fig. 5, we represent the response times (Fig. 5.A) and choice accuracy (Fig. 5.B) with respect to the reported confidence level for each participant. The data points show the experimental results (with the error bars as the bootstrapped 95% confidence interval), and the colored line the result of the simulation (with the light colored area the bootstrapped 95% confidence interval). Response times decrease Baranski and Petrusic [1994], Desender et al. [2018a] and accuracies increase with confidence Geller and Whitman [1973], Vickers and Packer [1982], Sanders et al. [2016], Desender et al. [2018a]. We find a monotonic dependency between response times and confidence, and between accuracy and confidence, but with specific shapes for each participant. Note that some values of confidence are only observed for a few trials, resulting then in large error bars especially for accuracy as we take the mean of a binary variable. For the numerical simulations, the relatively large size of the confidence interval is due to the limited number of trials, since we limit ourselves to the same protocol as the experimental one.

**Figure 5:**
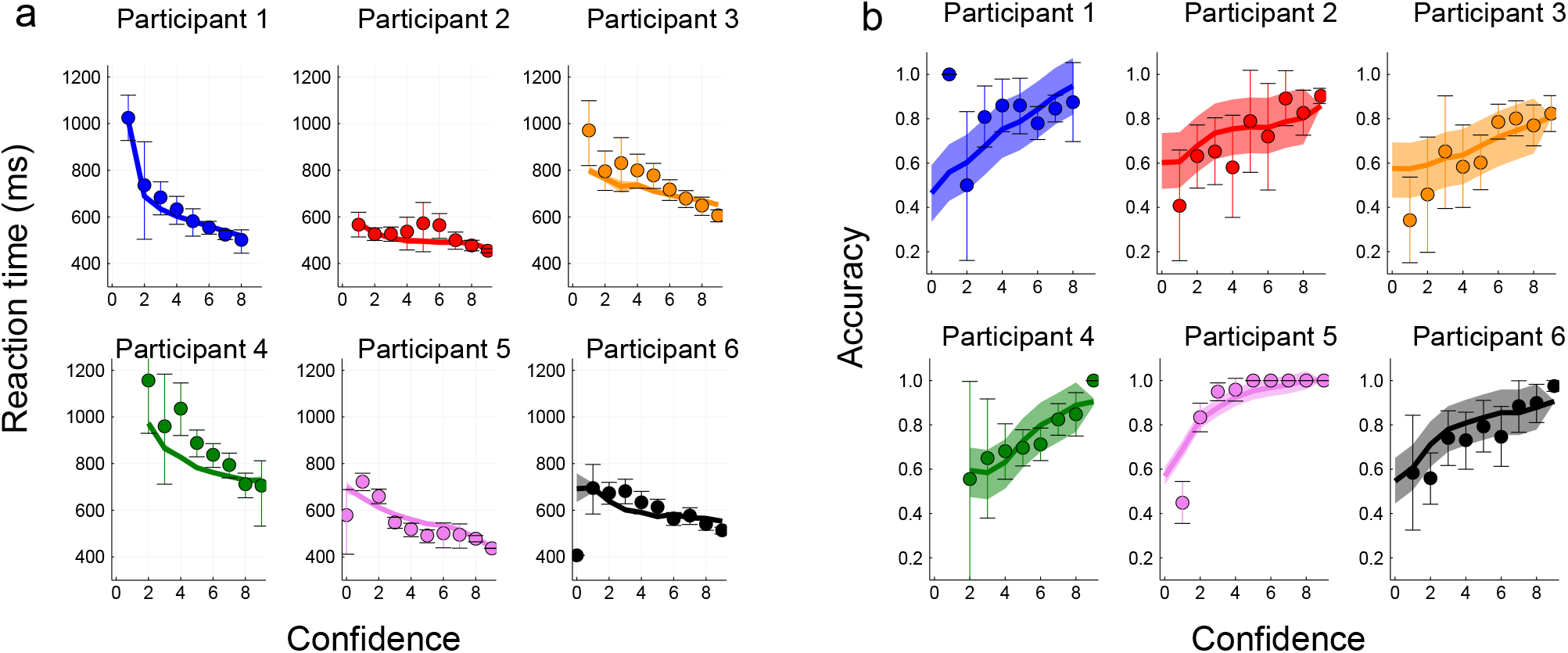
Response times and Accuracy as a function of confidence. (A) Response times, (B) Accuracy. For both panels: each sub-panel represents a different participant. Dots are experimental data with 95% bootstrapped confidence interval as error bars. Lines are averages over 20 simulations of the attractor neural network model (calibrated as explained in the Methods Section). The shaded area represents the 95% bootstrapped confidence interval on the mean.

For comparison, we also fit on our experimental data another non-linear model with mutual inhibition, the Usher-McClelland model Usher and McClelland [2001] (see Supplementary Information Section S1). The model fits the response times with respect to confidence, but only at intermediate levels of confidence. For some participants, we observe a strong divergence at high confidence (Participant 1,4 and 5). Although accuracy is an increasing function of confidence (except for participant 5), the experimental data do not fall within the bootstrapped confidence interval of the simulations (Supplementary Information Fig. S1). In contrast, we see that our more biophysical model correctly reproduces the psychometric and chronometric functions with respect to confidence for each participant, despite the important difference of response times between participants.

Previous studies found that, during a perceptual task, reported confidence increases with stimulus strength for correct trials, but decreases for error trials Kepecs et al. [2008], Sanders et al. [2016], Desender et al. [2018a]. This effect of confidence has been correlated to patterns of firing rates in experiments with rats Kepecs et al. [2008] and to the human feeling of confidence Sanders et al. [2016]. We observe the same type of variations of confidence with respect to stimulus strength (Supplementary Information Fig. S2), both in the experimental results and in the model simulations. This effect is in accordance with a prediction of statistical confidence, defined as the Bayesian posterior probability that the decision-maker is correct Griffin and Tversky [1992], Ernst and Banks [2002], Sanders et al. [2016]. We thus see that the attractor network model reproduces a key feature of statistical confidence.

## Impact of confidence on history biases

### Statistical analysis of sequential effects

Perceptual decisions made by humans in behavioral experiments depend not only on the current sensory input, but also on the choices made at previous trials. Various sequential effects have been reported Laming [1979], Leopold et al. [2002], Gold et al. [2008], and researchers have proposed different models to account for them Cho et al. [2002], Glaze et al. [2015], Bonaiuto et al. [2016], Berlemont and Nadal [2019] – in a previous work on post error effects Berlemont and Nadal [2019] we give a more general discussion of sequential effects. When the subject does not receive any feedback, confidence in his/her decision might be important for controlling future behaviors Yeung and Summerfield [2012], Meyniel et al. [2015b]. Recently, the effects of confidence on the history biases have been experimentally investigated Braun et al. [2018], Samaha et al. [2018]. One main finding is that decisions with high confidence confer stronger biases upon the following trials. Here, we investigate the influence of confidence upon the next trial in the empirical data, and we show that the results are well reproduced by the behavior of the dynamical neural model.

First, we perform a statistical analysis of the effect of history biases on response times in the experimental data.

For this, for each participant, we classify each trial into *low* and *high* confidence: a trial is considered as *low confidence* (resp. *high confidence*) if the reported confidence is below (resp. above) the participant’s median. We analyze the history biases making use of linear mixed effects models (LMM) Gelman and Hill [2007]. The LMM we consider assumes that the logarithm of the response time at trial *n*, *RT*_*n*_, is a linear combination of factors as follows:

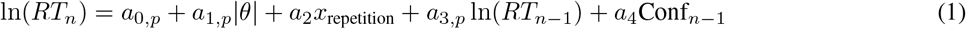

with *x*_repetition_ a binary variable taking the value 1 if the correct choice for the current trial is a repetition of the previous choice (and 0 otherwise), *θ* the orientation of the Gabor (in degree), *RT*_*n* − 1_ the response times of the previous trials, and Conf_*n*− 1_ the confidence of the previous trial coded as 0 for *low* and 1 for *high*. The subscript *p* in a coefficient (e.g *a*_0,*p*_) indicates that for this parameter we allow for a random slope per participant. We show the results in Table 1.

We find that higher orientations lead to faster response times and the repetition biases on response times Cho et al. [2002]. In line with previous works Desender et al. [2018a], high confidence has the effect of speeding up the following trial. Finally, we find that the previous response time has an effect on the subsequent one, meaning that the participants have the tendency to show sequences of fast (or slow) response times.

**Table 1:**
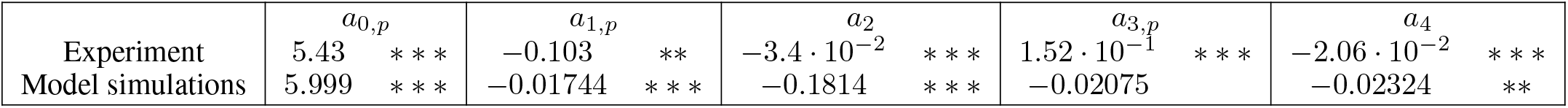
Results of the application of the LMM on the experimental data (first row) and on the data from the neural network simulations (second row). Each column correspond to a different parameter of the LMM, equation (1). We note for *p <* 0.005 and for *p* < 0.001. For more details on the statistical analysis, see Methods and the Supplementary Information, section S3.

Next, following a numerical protocol replicating the experimental one, for each participant we make numerical simulations with the model specifically calibrated on the participant’s data. Recall that the fit has been done on mean response times and accuracy, hence without taking into account serial dependencies. We then look at correlations between decisions made in successive trials by the neural attractor network, performing the same type of statistical analysis as done on the experimental data (see Methods, Table 1).

We note that the attractor neural network captures the variation of response times with respect to angle orientation Wong and Wang [2006]. We find that the dependency in the choice history (through the repetition of responses), as observed in the experimental data, is correctly reproduced by the model, in agreement with a previous study of these effects Berlemont and Nadal [2019]. Quite remarkably, we observe an effect of confidence on response times in the network, with the same sense of variation (negative slopes) as in the experiment.

### Analysis of the underlying neural dynamics

To understand how the neural dynamics leads to these confidence-specific sequential effects, we make an analysis of the dynamics similar to the one done in Berlemont and Nadal Berlemont and Nadal [2019] for the analysis of post-error effects in the same neural model. We illustrate this analysis in Fig. 6. On each panel, we compare the *mean* neural dynamics for post-low and post-high confidence trials (respectively red and blue lines). Without loss of generality, we assume that the previous decision was a *C* grating. We first note that the relaxation dynamics between two consecutive trials are different, resulting in different starting points for the next trial, from post-low and post-high confidence trials. Panel (A) corresponds to the case where the new stimulus is also *C* oriented ("repeated" case), at low strength level. The ending points of the relaxations fall into the correct basin of attraction. Because the post-high confidence relaxation lies deeper into the basin of attraction than the one of post-low trials, the subsequent dynamics will be faster for post-high confidence trials in this case. In panel (B) we represent the case, still at low stimulus strength, where the stimulus orientation of the new stimulus is the opposite ("alternated" case) to the one corresponding to the previous decision (hence an *AC* grating). Both dynamics lie close to the basin boundary of the two attractors, thus the dynamics are slow and there is no significant difference between post-low and post-high confidence trials. In panels (C) and (D) we represent the same situations as panels (A) and (B), respectively, but for high strength levels (easy trials). The ending points of the relaxations are far from the boundary of the basins of attraction, whatever the grating presented. The response times for post-high and post-low confidence trials are thus similar. This analysis shows that the non-linearity of the network dynamics is responsible for the considered sequential effect. More precisely, it is the very existence of basin boundaries, and the fact that the network state is more or less close to the basin boundary at the onset of a new stimulus, which lead to the sequential effects.

**Figure 6:**
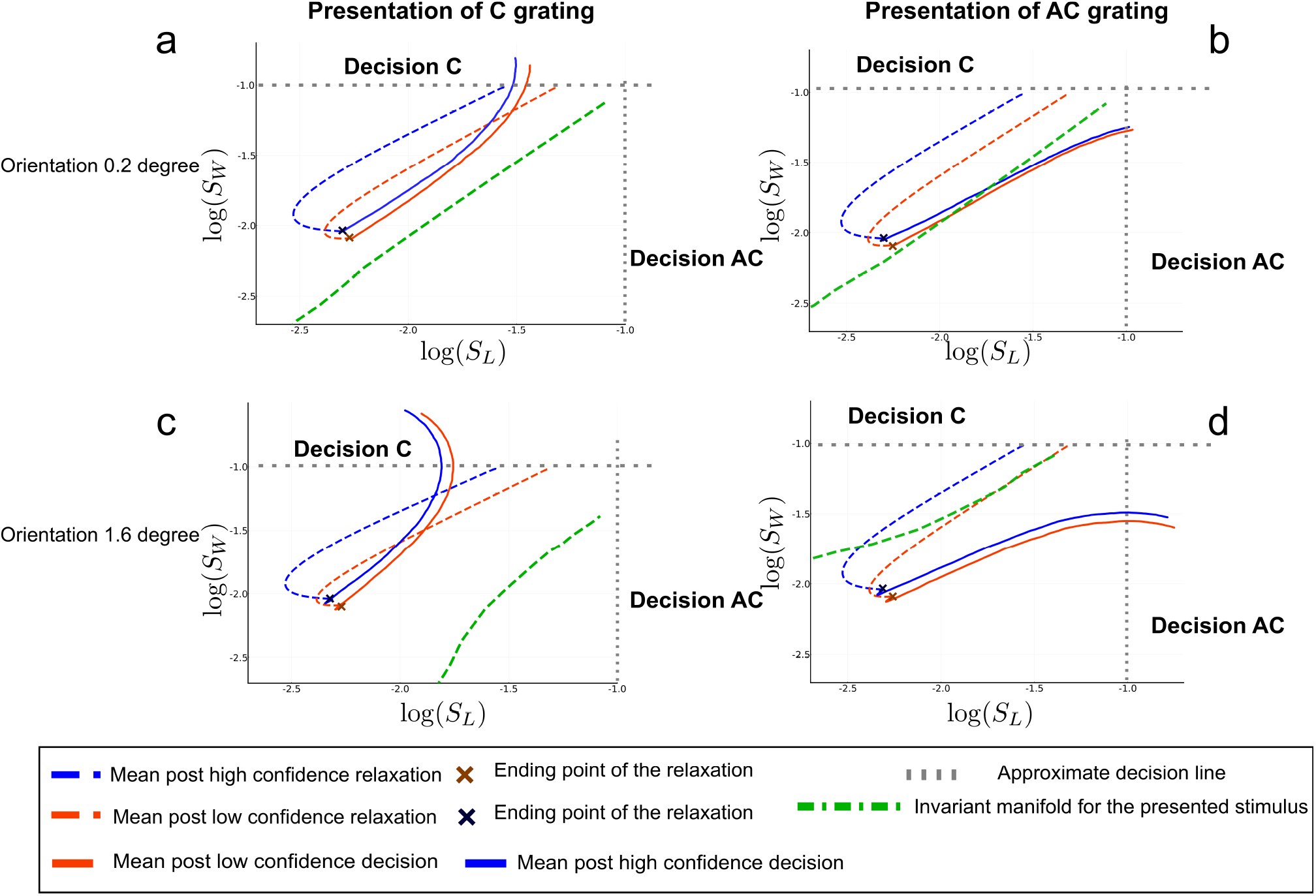
Non linear dynamics in post-low and post-high confidence trials. Phase-plane trajectories (in log-log plot, for ease of viewing) of the post low and high confidence trials. We assume that the previous decision was decision C. The axis represent the losing neural pool S_L_ and and the winning neural pool S_W_ at the previous trial. The blue color codes for post-high confidence trials, and the red one for post-low confidence. Panels (A) and (B): Repeated and alternated case for low orientation stimuli; Panels (C) and (D): Repeated and alternated case for high orientation stimuli. In order to compare the decision times, the dynamics starting at the onset of the next stimulus is followed during 200ms, as if there were no decision threshold. The actual decision occurs at the crossing of the dashed gray line, indicating the threshold.

We now qualitatively confront the outcomes of the above analysis with the experimental data. To do so, we group the response times according to the same cases as previously: high and low stimulus strength, repeated or alternated trials. We compare post high and low confidence trials in each case, using a t-test Fay and Proschan [2010]. We find that mean response times between post low and high confidence trials are different in the low orientation stimuli and repeated case (t-test, *p* = 0.044, *df* = 1322.6), but they are identical in the high orientation stimuli and alternated case, high orientation stimuli and alternated case, low orientation stimuli and repeated case they are identical (respectively *p* = 0.90, *df* = 778.7, *p* = 0.70, *df* = 610, *p* = 0.23, *df* = 617.4). This is in accordance with the outcomes of the above analysis based on the non-linear dynamics.

The model reproduces sequential effects correlated with repetition and confidence, and we have shown that these effects result from the intrinsic nonlinear network dynamics. However, the model does not induce correlations between response times of two successive trials (for more details see Supplementary Information S3). This suggests that the correlations observed in the experimental data cannot be explained by the intrinsic dynamics of the attractor network, but may come from higher order processing. Within a DDM approach, one may account for such effect, but only with a change of parameters from trial to trial (e.g. by changing the decision threshold depending on the previous reaction time). However, such ad hoc changes of parameter values are not supported by models or experimental data which would provide clues about the neurodynamical mechanisms underlying these changes.

### Comparison with diffusion models

Next we investigate if linear models of decision-making of the DDM family are able to reproduce the sequential effects. More precisely, we consider the independent race model (IRM Raab [1962], Vickers [1970], Merkle and Van Zandt [2006]). During the accumulation of evidence the equations of evolution of the activities *x*_*i*_, *i* = *{C, AC}*, are:

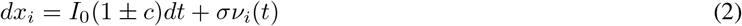

where *ν*_*i*_(*t*), *i* = {*C, AC*}, are white noise processes. The first race that reaches a threshold *z* (or −*z*) is the winning race. The confidence in the decision is modelled as a monotonic function of the balance of evidence |*z* − *x*_*losing*_| Vickers [1979 (reeditited in 2014], Drugowitsch et al. [2014], Mamassian [2015].

We extend the IRM in order to deal with sequences of trials. To do so, we allow for a relaxation dynamics between trials, in a way analogous to the relaxation dynamics in the attractor network model. Hence, after a decision is made, both units receive a non specific inhibitory input leading to a relaxation until the next stimulus is presented (see Fig. 7). Within this extended IRM framework, we study how, *with a fixed set of parameter values*, the sequential effects would be correlated with confidence.

**Figure 7:**
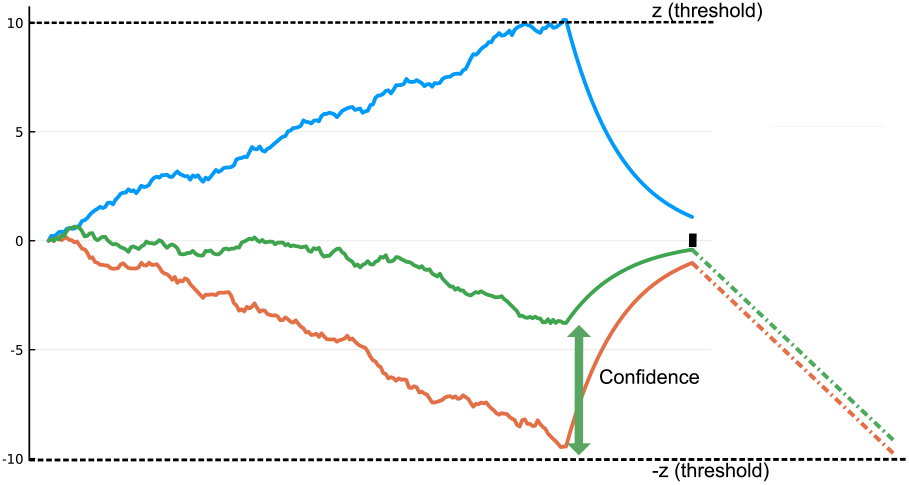
Schematic dynamics of a race model with a relaxation mechanism. The upper and bottom dash lines correspond to the two opposite decision thresholds. The blue trajectory is a typical winning race. The black rectangle on the x-axis denotes the onset of the next stimulus, hence the end of the relaxation period. The green and orange trajectories are the loosing races in two trials with different confidence outcomes. The green and orange dashed lines represent the mean dynamics of these two races during the presentation of the next stimulus.

Since in the IRM there is no interaction between the two races, the relaxation of the winning race is the same in both low and high confidence trials. However, the ending point of the relaxation following a decision is closer to the base-line (0 line) for a high confidence trial than when it comes to a trial with low confidence trial (Fig. 7). For the next trial, if the winning race is the same as previously, then the mean response times are identical in low and high confidence cases. However, if the opposite decision is made, the response time in the post-low confidence case is faster than the one in the post-high confidence case, as we can observe with the mean race shown in Fig. 7. This behavior is in contradiction with the experimental data for which we observe the opposite effect (see Supplementary Information Table S3). This conclusion applies more generally to any race-type model without interactions between units. This applies as well to IRMs with a non-linearity under the form of collapsing bounds (see Supplementary Information S6). Other researchers have studied different kind of sequential effects, such as post-error slowing, with collapsing bounds models Purcell and Kiani [2016]. However, these models do not have a relaxation dynamics between trials, and the model parameters are allowed to be modified between trials. Here we show that the confidence-specific sequential effects result from a type of non-linearity in the dynamics which impacts the interaction between neural units.

Authors have proposed a set of *non-linear* diffusion equations to approximate the process of decision-making in attractor neural networksRoxin and Ledberg [2008]. These equations result from a description of the dynamics at the vicinity of the bifurcation point - the particular state at which any small input will induce the appearance of the two attractors. This framework is not well adapted to model sequences of decisions, since one would need to reset the system at the vicinity of the bifurcation point before the onset of a new trial. Even if this would be done, the system would not account for the experimentally observed sequential effects, for the very same reasons as those presented above for linear diffusion models. As shown by our analysis of the attractor network dynamics, the key non-linearity underpinning the sequential effects is the one resulting from the existence of frontiers between basins of attractions.

## Discussion

Dynamical models of decision making implement in different ways the same qualitative idea: decision between two categories is based on the competition between units collecting evidences in favor of one or the other category (or with a single unit whose activity represent the difference between the categorical evidences). Most authors propose that behavioral confidence can be modeled as a function of the balance of evidence Vickers [1979 (reeditited in 2014], Kepecs et al. [2008], Moreno-Bote [2010], Pleskac and Busemeyer [2010], Zylberberg et al. [2012]. Very few works propose other mechanisms Rolls et al. [2010a,b], Paz et al. [2016]. Among these exceptions, the consensus model Paz et al. [2016] assumes that several attractor neural networks are run in parallel, and a decision is reached when there is a consensus (more than half of the networks have chosen the same alternative). Within this framework, confidence is defined as the fraction of networks that have chosen the winning decision. This model can account for the relation between confidence and reaction times Paz et al. [2016]. However, it is not clear how it can be modified in order to perform sequences of decisions. In particular, since the different networks reach the decision threshold at different times, it is not obvious how to decide when relaxation should start for each network, and it is not clear what kind of sequential effects should be expected for any given relaxation rule.

Considering the large majority of models based on the DDM framework, in view of our findings we now contrast the DDM and attractor neural network approaches. We do so on three aspects: the modelling of confidence, the analysis of sequential effects, and the issue of non-decision times.

Bayesian inference models compute confidence using drift-diffusion models (DDM) extensions based on decision variable balance Vickers [1979 (reeditited in 2014], Kepecs et al. [2008], Moreno-Bote [2010], Zylberberg et al. [2012], possibly with additional mechanisms - decision variable balance combined with response times Kiani et al. [2014] or post-decisional deliberation Pleskac and Busemeyer [2010] (the dynamics continues after the decision, thus updating the balance of evidence). Similar studies have been made with independent race models (IRM) Raab [1962], Vickers [1970], Merkle and Van Zandt [2006]. These DDM or IRM models successfully account for various psychometric and chronometric specificities of human confidence. In DDMs, confidence based on decision variable balance predicts that confidence should deterministically decrease as a function of response times Kiani and Shadlen [2009]. However, the response times distributions strongly overlap across confidence levels Ratcliff and Starns [2009]. This property can be recovered making use of additional processes, such as with a two-stage drift-diffusion model Pleskac and Busemeyer [2010]. Yet, other effects remain unexplained within the framework of DDM. This is the case of early influence of sensory evidence on confidence Zylberberg et al. [2012], as well as the fact that confidence is mainly influenced by evidence in favor of the selected choice Zylberberg et al. [2012].

Within the framework of attractor neural networks, early sensory evidence influence decision accuracy and reaction time Wong et al. [2007]. The model discussed in the present paper is appropriate for going beyond and studying the effect of early sensory evidence on confidence. Given our findings, we expect that the model will reproduce the above mentioned effects not well accounted for by the DDMs.

As discussed in the Results section, various serial dependence effects are observed in perceptual decision-making. A recent finding is that the magnitude of history biases increases when previous trials were faster and correct. Given the known correlations between confidence, response time and accuracy, this effect is interpreted as an impact of confidence on the next decision Braun et al. [2018]. By measuring directly the subjective confidence of the participants, recent studies confirm that history biases are correlated with confidence Desender et al. [2018a], Samaha et al. [2018], Desender et al. [2018b]. In our experiment, we observe that high confidence trials lead to faster subsequent choices in agreement with the above mentioned experimental studies. On the theoretical side, this impact of confidence on response times of the subsequent trials has been investigated within the framework of DDMs Desender et al. [2018a]. The usual analysis consists in dividing trials into two categories, subsequent to low or high confidence trials, fitting then separately a DDM on each type of trial. The main finding is that the parameters (threshold and drift) are different depending on the confidence level at the previous trial. In the absence of changes of parameters, even with the addition of a relaxation between trials as discussed in the Results section, the DDM or IRM models cannot account for the observed sequential effects: as we have seen, the predicted sequential effect would be the opposite of the observed one. In contrast, as discussed in the Results section, with a unique choice of parameters values for each participant, the attractor network model does not only account for the relationship between confidence, response times and accuracy, but also reproduces the influence of confidence on serial dependence.

Finally, the question of the non-decision times arise from our modelling work. Human studies commonly report right-skewed response times distribution Luce et al. [1986], Ratcliff and Rouder [1998]. Such long right tails are well captured by drift-diffusion models Ratcliff and Rouder [1998], Usher and McClelland [2001] - and this is generally considered as a strong evidence in favor of the accumulation of evidence mechanism. However, with trained subjects, the right-skew is less pronounced and a Gaussian distribution fits well the response times distribution Peirce [1873]. In contrast to human studies, experiments in monkeys do not show such long right tails in the response times histograms Ditterich [2006]. When assuming a constant value for the non-decision time, attractor neural network models do not produce right-skewed distributions, but accurately reproduce the shape of the distributions in monkeys experiments Wang [2008]. In accordance with these results, in this work we have shown that for the range of parameters we considered, the decision time distribution generated by the neural network can be approximated by a Gaussian distribution. Within the neural attractor framework, the experimentally observed long right tails can thus be understood as originating only from the non-decision times. Here we have proposed an estimation of the distribution of these non-decision time allowing to fit the empirical response times distributions. One should note that, even in the case of an analysis of experimental data within the DDM framework, the estimated non-decision times are not necessarily given by a constant value, but may show a distribution with a strong right skew Verdonck and Tuerlinckx [2016]. These findings combined with ours suggest that the question of the origin of the long right tails in human response times has to be reconsidered.

To conclude, in this work we designed an experiment in order to study confidence with human participants. We fitted a neural attractor network model specifically to each participant in order to describe their behavioral results in continuous sequences of perceptual decisions: response times, accuracy and confidence. Quite remarkably, we found that the impact of confidence on sequential effects is well described by the nonlinear nature of the attractor dynamics.

## Methods

### Experiment

#### Participants

Nine participants (7 Females, Mean Age = 27.3, SD = 5.14) have been recruited from the Laboratoire de Psychologie cognitive et de Psycholinguistique’s database (LSCP, DEC, ENS-EHESS-CNRS, PSL, Paris, France). Every subject had normal or corrected-to-normal vision. The participants performed three sessions on three distinct days in the same week for a total duration of about 2h15. Three participants were excluded. Two of the excluded participants did not complete correctly the experiment and one exhibited substantially asymmetric performance (98% of correct responses for an angle of 0.2°, but 18% at −0.2°degree). As a result, we analyzed data from 6 participants.

#### Ethics statement

The experiment followed the ethics requirements of the Declaration of Helsinki (2008) and has been approved by the local Ethics Committee (Comité de Protection des Personnes, Ile-de-France VI, Paris, France). We obtained written informed consent from every participant who received a compensation of 15 euros for their participation.

#### Stimuli and tasks

The stimuli were generated using Matlab along with the Psychophysics Toolbox Kleiner et al. [2007]. They were displayed on a monitor at 57.3 cm of the participants’ head. The participants performed the experiment in a quiet and darkened experimental room. Their heads were stabilized thanks to a chin-rest. Trials began with the presentation of a black fixation point (duration = 200 ms). Then the stimulus for the primary decision task was presented, consisting in a circular grating (diameter = 4°, Tukey window, 2 cycles per degree, Michelson contrast = 89%, duration = 100 ms, phase randomly selected at each trial). The grating had eight possible orientations with respect to the vertical meridian, and participants were asked to categorize them as clockwise or anti-clockwise with respect to the vertical meridian by pressing the right-arrow or left-arrow. Participants had been instructed to respond as follows: “You have to respond quickly but not at the expense of precision. After 1.5 s the message, "Please answer", will appear on the screen. It would be really ideal, if you would answer before this message appears.”.

Trials were of three types, grouped in *pure* block, *feedback* block and *confidence* block (see below). Participants performed three sessions on three distinct days. Each session (45 min) consisted in three runs, each run being composed of one exemplar of each of the three type of block, in a random order. Before starting the experiment, participants performed a short training block of each type, with easier orientations than in the main experiment.

#### Pure block

In this block, participants waited 300 ms after each decision, before the black fixation point appears. The stimulus appeared 200 ms after this fixation point. The eight possible orientations for the circular grating were [-1.6°, −0.8°, −0.5°, −0.2°, 0.2°, 0.5°, 0.8°, 1.6°] and a stimulus was chosen randomly among them with the following weights: [0.05, 0.1, 0.15, 0.2, 0.2, 0.15, 0.1, 0.05].

#### Feedback block

In this block, 200 ms after the decision, the participants received an auditory feedback (during 200 ms) about the correctness of the decision they just made. The black fixation dot appeared 100 ms after this feedback and a new trial began. The orientations of the circular gratings were chosen randomly from [-1.6°, −0.8°, −0.2°, 0.2°, 0.8°, 1.6°] with the following weights [0.12, 0.18, 0.2, 0.2, 0.18, 0.12].

#### Confidence block

In the confidence block, participants had to evaluate the confidence on the orientation task 200 ms after the decision. To perform this task they had to move a slider on a 10-points scale, from *pure guessing* to *certain to be correct*. Importantly, the initial position of the slider was chosen randomly for each trial. Participants moved the slider to the left by pressing the "q" key, and to the right with the "e" key. We ask the following kind of confidence judgment to the participants: one extreme of the scale is "pure guess", the other is "absolutely certain". They confirmed the choice of the value of confidence by pressing the space bar. The participants had the choice to indicate that they had made a "motor mistake" during the orientation task. For this they had to press a key with a red sticker instead of responding on the confidence scale. After the choice of confidence, the participants had to wait 300 ms before the black fixation dot appears. After the fixation dot the stimulus appeared 200 ms later. The orientations of the circular gratings were the same as in the feedback block.

Accuracy is higher and response times are slower in confidence blocks than in pure blocks. To test this effect on accuracy we ran a binomial regression of responses with fixed factors of orientation and type of block (pure or confidence), the interaction between these factors and a random participant intercept. The orientation coefficient was 2.15 (SD = 0.17, *z* = 12.44 and *p* < 10^−16^); there was no effect of block type (*p* = 0.385). But we found a significant orientation by block type interaction (value of 0.55, SD = 0.08, *z* = 6.97 and *p* = 3 ⋅ 10^−12^), indicating that participants were more accurate in confidence blocks than in no-confidence blocks. In a similar way, we test the effect on response times by using a mixed effect regression with the same factors and intercept as for the accuracy (only on the absolute value of the orientation). We found that the orientation coefficient (value of −0.08, SD = 0.013 and *p* = 0.0006) and the block type coefficient (value of 0.095, SD = 0.028 and *p* = 0.011) were significant, meaning that participant are slower in the confidence block. Moreover, the slope by block type interaction with orientation was also significant (value of −0.028, SD = 0.010 and *p* = 0.031), meaning that the difference between the two types of blocks is more important at low orientation. Surprisingly, we find that performance and response time across participants are identical in the feedback and pure blocks (no statistically significant difference). The participants were highly trained in the orientation discrimination task.

### Statistical Analyses

We used RStudio with the package *lme4* Bates et al. [2015] to perform a linear mixed effects analysis Gelman and Hill [2007] of the history biases of the reaction times, on both the experimental and the numerical simulations data.

To perform the comparison between the experimental data and the model results illustrated in Fig. 6, we first transform the response times of each participant using the z-score Kreyszig [1979]. This normalization allows analyzing the participants all together.

We compared the LMM model described in the main text, equation (1), to other ones that do not include all the terms, using the *ANOVA* function (with the lme4 package Bates et al. [2015]) that performs model comparison based on the Akaike and Bayesian Information Criteria (AIC and BIC) Bates et al. [2015]. We find that the LMM from equation (1) is preferable in all cases (see Supplementary Information Table S4).

### Attractor neural network model

#### Neural dynamics

We consider a decision-making recurrent network model governed by local excitation and feedback inhibition Wong and Wang [2006], Berlemont and Nadal [2019]. Within a mean-field approach, Wang and Wong Wong and Wang [2006] have derived a reduced firing-rate model of a detailed biophysical model of spiking neurons Compte et al. [2000]. This reduced model is composed of two interacting neural pools which faithfully reproduces not only the behavioral behavior of the full model, but also the dynamics of the neural firing rates and of the output synaptic gating variables. The model variant Berlemont and Nadal [2019] that we consider here takes into account a corollary discharge Sommer and Wurtz [2008], Crapse and Sommer [2009]. This results in a non specific inhibitory current injected into the neural pools just after a decision is made, making the neural activities relax towards a low activity, neutral, state, therefore allowing the network to deal with consecutive sequences of decision making trials Berlemont and Nadal [2019]. For completeness, we recall here the equations and parameters with notation adapted to the present study.

The model consists of two competing units, each one representing an excitatory neuronal pool, selective to one of the two categories, *C* or *AC*. The dynamics is described by a set of coupled equations for the synaptic activities *S*_*C*_ and *S*_*AC*_ of the two units *C* and *AC*:

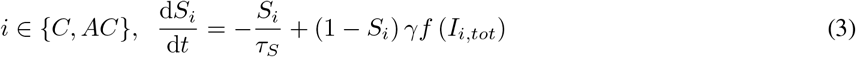

The synaptic drive *S*_*i*_ for pool *i* ∈ {*C, AC*} corresponds to the fraction of activated NMDA conductance, and *I*_*i,tot*_ is the total synaptic input current to unit *i*. The function *f* is the effective single-cell input-output relation Abbott and Chance [2005], giving the firing rate as a function of the input current:

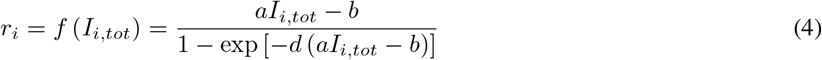

where *a, b, d* are parameters whose values are obtained through numerical fit. The total synaptic input currents, taking into account the inhibition between populations, the self-excitation, the background current and the stimulus-selective current, can be written as:

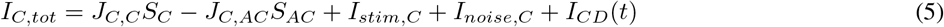

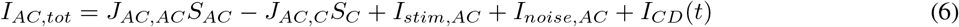

with *J*_*i,j*_ the synaptic couplings. The minus signs in the equations make explicit the fact that the inter-units connections are inhibitory (the synaptic parameters *J*_*i,j*_ being thus positive or null). The term *I*_*stim,i*_ is the stimulus-selective external input. The form of this stimulus-selective current is:

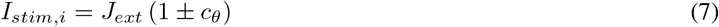

with *i* = *C, AC*. The sign, ±, is positive when the stimulus favors population *C*, negative in the other case. Here the parameter *J*_*ext*_ combines a synaptic coupling variable and the global strength of the signal (which are parametrized separately in the original model Wong and Wang [2006], Berlemont and Nadal [2019]). The quantity *c*_*θ*_, between 0 and 1, characterizes the stimulus strength in favor of the actual category, here an increasing function of the (absolute value of) the stimulus orientation angle, *θ*.

In addition to the stimulus-selective part, each unit receives individually an extra noisy input, fluctuating around the mean effective external input *I*_0_:

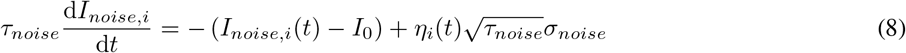

with *τ*_*noise*_ a synaptic time constant which filter the white-noise.

On presentation of a stimulus, the system evolves toward one of the two attractor states, corresponding to the decision state. We consider that the decision is made when for the first time the firing rate of one of the two units crosses a threshold *z*.

After each decision, a corollary discharge under the form of an inhibitory input is sent to both units until the next stimulus is presented:

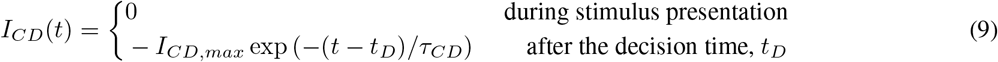

This inhibitory input, delivered between the time of decision and the presentation of the next stimulus, allows the network to escape from the current attractor and thus engage in a new decision task Berlemont and Nadal [2019].

#### Confidence modeling

Within the various decision making modelling frameworks, similar proposals have been made to model the neural correlated of the behavioral confidence level. In race models Raab [1962], which have equal number of accumulation variables and stimulus categories, as in attractor network models, the balance of evidence at the time of perceptual decisions has been used to model the neural correlate of the behavioral confidence Vickers and Packer [1982], Smith and Vickers [1988], Wei and Wang [2015]. This balance of evidence is given by the absolute difference between the activities of the category specific units at the time of decision. Here, we consider that confidence is obtained as a function *f* of the difference in neural pools activities Wei and Wang [2015], ∆*r* = |*r*_*C*_ − *r*_*AC*_|.

In our experiment, the subjects expressed their confidence level by a number on a scale from 0 to 9. In order to match the neural balance of evidence with the confidence reported by the subject, we map the balance of evidence histogram onto the behavioral confidence histogram, a procedure called *histogram matching* Gonzalez et al. [2002]. Note that the mapping is here from a continuous variable to a discrete one (taking integer values from 0 to 9).

### Fitting procedure

For each participant we calibrate the model by fitting both the mean response times and the accuracies for each orientation, this separately for each block. We note that we only fit the means, which in particular implies that the fits do not take into account the serial dependencies. Doing so, any sequential effects that will arise in the model will result from the intrinsic dynamics of the network, and not from a fitting procedure of these effects.

For most model parameters we take the value used in a previous study Berlemont and Nadal [2019], as reproduced in Table 2. For the models calibration we consider *I*_*CD,max*_, *τ*_*CD*_, *c*_*θ*_ and *z* as free parameters. We impose the two parameters *I*_*CD,max*_ and *τ*_*CD*_ to be common to all participants (joint optimization). We optimize the parameters *c*_*θ*_ (one for each orientation value) and the decision threshold *z* across subjects and blocks.

**Table 2:** Numerical values of the model parameters. The left and middle panel are parameters common to all participants Berlemont and Nadal [2019]. The right panel lists the parameters which are fitted on the behavioral data. The paramerer *c*_*θ*_ is a function of *θ* that is either linear or quadratic.

| Parameter | Value | Parameter | Value | Fitted parameter | Status |
| --- | --- | --- | --- | --- | --- |
| a | 270 Hz/nA | $\sigma_{noise}$ | 0.02 nA | $I_{CD,max}$ | common to all participants |
| b | 108 Hz | $\tau_{noise}$ | 2 mS | $\tau_{CD}$ | common to all participants |
| d | 0.154 s | $I_0$ | 0.3255 nA | decision threshold $z$ | specific to each participant |
| $\gamma$ | 0.641 | $J_{ext}$ | 0.182 nA | $c_\theta = \alpha_1 \theta + \alpha_2 \theta^2$ | specific to each participant |
| $J_{C,C} = J_{AC,AC}$ | 0.2609 nA | $J_{C,AC} = J_{AC,C}$ | 0.0497 nA | | |
| $\tau_S$ | 100 ms | | | | |

The rationale for this choice of free parameters is as follows. To avoid overfit, one has to restrict as much as possible the number of free parameters. We rely on the model calibrations done in previous works Wang [2002], Wong and Wang [2006], Berlemont and Nadal [2019] which suggest to keep as much as possible the parameters values resulting from the initial work of Wong and Wang. In particular, the original parameters values were chosen such as to reproduce empirical data with the mean field model. Now since the empirical data are only behavioral data, it is difficult to make a calibration of the synaptic weights. A significant change of these parameters would be required to change the behavioral outcomes. Importantly, we tried to restrict the calibration to a small set of reasonably independent parameters. For instance, a change in the weights values may be compensated by a change in the decision threshold (so that the cost function may be flat on a large domain of the parameters space). With the weights fixed, we can optimize the fit with respect to the decision threshold in a safer way. An important quantity is the signal-to-noise ratio. By keeping the internal noise constant during the fitting procedure, we explore the whole range of this ratio. We also note that our choice of free parameters can be paralleled with the one made in the DDM framework (drift, threshold and level of noise). This facilitates the comparison with the DDM approach. Finally, imposing some of the free parameters to be common to all participants allows us to further reduce the number of free parameters, at a price of a more complex optimization (a partially joint calibration of all the participant-specific networks).

The observed response time is the sum of a decision time and of a non decision time. Assuming no correlation between these two times, the mean non decision time is thus independent of the orientation. For comparing data with model simulations (which only gives a decision time) at any given orientation *θ*, we first substract to the mean response time the mean response time averaged over all orientations (this for both data and simulations). We calibrate the model parameters so as to fit these centered mean response times. This will provide a fit of the mean response times (at each angle) up to a global constant, which is the mean non decision time (the modeling of the non decision time distribution is presented in the next Section).

For each participant, and each block, we thus consider the cost function:

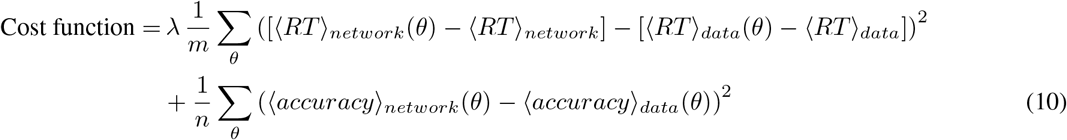

where the sums are over the orientation values, *θ* = {0.2°, 0.5°, 0.8°, 1.6°, the brackets 〈…〉 design averages (as detailed below), and the normalization factors *n* (for response times) and *m* (for the accuracy) are given by

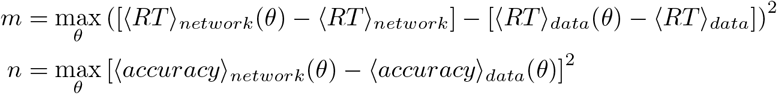

In these expression, 〈*RT*〉_*data*_(*θ*) denotes the mean experimental response time obtained by averaging over all trials at the orientations ±*θ*, 〈*RT*〉_*data*_ is the average over all orientations; 〈*RT*〉_*network*_(*θ*) and 〈*RT*〉_*network*_ are the corresponding averages obtained from the model simulations. The coefficient *λ* denotes the relative weight given to the response time and accuracy cost terms. We present the results obtained when taking *λ* = 2, but it should be noted that the choice of this parameter does not impact drastically the fitted parameters.

For each subject, we minimize this cost function with respect to the choice of *c*_*θ*_ and *z*, making use of a Monte Carlo Markov Chain fitting procedure, coupled to a subplex procedure Rowan [1990]. This method is particularly adapted to handle simulation based models with stochastic dynamics. Finally, *I*_*CD,max*_ and *τ*_*CD*_ are fitted using a grid search algorithm as they have less influence on the cost function. In the model, the parameter *c* represents the stimulus ambiguity, which we expect here to be a monotonous function of the amplitude of the angle, *θ*. When allowed to be independent parameter values for each value of the orientation, *θ* = 0.2°, 0.5°, 0.8°, 1.6°, we find that the *c*_*θ*_ values can be approximated by a linear or quadratic function of *θ* depending on the participant. We performed an AIC test Akaike [1992] between the linear and quadratic fit in order to choose which function to use for each participant. These approximations reduce the number of free parameters.

In order to obtain a confidence interval for the different parameters, we used the likelihood estimation of confidence interval for Monte-Carlo Markov Chains method. The confidence interval on the parameters is thus the 70% confidence interval, assuming a Gaussian distribution of the cost function. This provides an approximation of the reliability of the parameters values found. In order to assess the reliability of this method we checked that the threshold *z* and stimulus strength *c*_*θ*_ parameters have an almost non-correlated influence onto the cost function. The results of the calibrating procedure are summarized in Supplementary Information Tables S5 and S6, with *I*_*CD,max*_ = 0.033 nA and *τ*_*CD*_ = 150 ms.

### Estimating the non-decision time

The above fitting procedure calibrates the mean response times up to a global constant, corresponding to the mean non decision time. As explained in the main text, we can go beyond and actually model the non-decision time distribution.

The non-decision time is considered to be due to encoding and motor execution Luce et al. [1986]. Most model-based data analysis of response time distributions assume a constant non-decision time Ratcliff and Rouder [1998], Usher and McClelland [2001], Wong and Wang [2006]. However, fitting data originating from a skewed distribution under the assumption of a nonskewed non-decision time distribution is cause for bias in the parameter estimates if the model for non-decision time is not correct Ratcliff [2013]. Recently, authors have proposed a mathematical method to fit a nonparametrical non-decision time Verdonck and Tuerlinckx [2016]. Analyzing various experimental data with this method within the framework of drift-diffusion models, they find that strongly right skewed non-decision time distributions are common.

In this paper we make the hypothesis that the non-decision time distributions are ex-Gaussian distributions, whose parameters are inferred from the data making use of the deconvolution method Verdonck and Tuerlinckx [2016] and detailed in Supplementary Information Section S5. We present in Fig. 3 the fits of the response time distributions and the inferred non decision time distributions.

## Acknowledgements

We are grateful to Laurent Bonnasse-Gahot for useful discussions and suggestions. We thank Pascal Mamassian, Vincent de Gardelle and Xiao-Jing Wang for stimulating discussions. We thank Isabelle Brunet for her help in recruiting the participants and organizing the experimental sessions. We thank the anonymous referees for useful remarks. KB acknowledges a fellowship from the ENS Paris-Saclay.

## Author contributions statement

All authors contributed to the research plan. K.B., J.-R.M. and J.S. conceived the experiment; K.B. and J.-R.M. conducted the experiment; K.B. and J.S. performed the statistical analyses; K.B. and J.-P.N. performed the mathematical modeling; K. B. performed the numerical simulations; K.B. and J.-P.N. analyzed the results with inputs from the other authors. K.B. and J.-P.N. wrote the paper with input from the other authors. All authors reviewed the manuscript.

### Additional information

#### Data and Source codes Availability

Experimental Data as well as the statistical analyses scripts are available at https://osf.io/eh2xb/. Simulations of the model were carried out using the Julia programming language Bezanson et al. [2017]. Scripts corresponding to these numerical simulations are also available at https://osf.io/eh2xb/.

#### Competing interests

The authors declare no competing interests.

