## Supplementary Information for "NONLINEAR NEURAL NETWORK DYNAMICS ACCOUNTS FOR HUMAN CONFIDENCE IN A SEQUENCE OF PERCEPTUAL DECISIONS"

#### S1 - Usher and McClelland model

We compare the fit with the attractor neural network with the fit with the Usher and McClelland model [Usher and McClelland, 2001]. The equations of the model are the following:

$$\tau dx_1 = -kx_1 dt - \beta f(x_2)dt + I_1 + \sigma\mu_1(t)$$
$$\tau dx_2 = -kx_2 dt + \beta f(x_1)dt + I_2 + \sigma\mu_2(t)$$

with $\mu_i(t)$ a white-noise process and $I_i$ the input current to the system. The external input is defined as $I_i = 0.5 \pm c_\theta$, with $c_\theta$ the strength per angle as in the attractor neural network. $\sigma = 0.4$ denotes the strength of the noise, $k$ the relaxation strength, $\tau = 0.1$ the relaxation time and $\beta$ the inhibitory term. Finally, the function $f$ is a sigmoidal function of gain $G = 0.4$ and half-activity offset $d = 0.5$, $f(x_i) = 1/\left[1 + \exp\left(-G(x_i - d)\right)\right]$. The dynamics occurs until a threshold $z$ is reached for one of the two units.

It should be noted that, despite the non-linearity, the Usher-McClelland model is closer to drift-diffusion models than to biophysical attractor model (this because the only non-linearity is in the interaction between units). Reductions to one-dimensional drift diffusion models can be made in various ranges of parameters Bogacz et al. [2006].

In order to fit this model to the experimental data, we apply the same procedure as for our attractor network model.

| **Parameter** | **Subject 1** | **Subject 2** | **Subject 3** | **Subject 4** | **Subject 5** | **Subject 6** |
|---|---|---|---|---|---|---|
| $z$ | 1.0 | 1.0 | 1.0 | 1.4 | 1.3 | 1.3 |
| $\beta$ | 0.25 | 0.10 | 0.18 | 0.10 | 0.15 | 0.12 |
| $k$ | 0.15 | 0.18 | 0.18 | 0.11 | 0.11 | 0.14 |
| $c_{0.2}$ | 0.02 | 0.04 | 0.02 | 0.02 | 0.02 | 0.04 |
| $c_{0.8}$ | 0.15 | 0.12 | 0.07 | 0.08 | 0.14 | 0.132 |
| $c_{1.6}$ | 0.23 | 0.20 | 0.17 | 0.225 | 0.295 | 0.235 |

Table S1: **Calibration of the Usher-McClelland model: Parameters for each participant after fit of the mean accuracy and response times of the *confidence* block.**

We note that the Usher-McClelland model fits the response times with respect to confidence, but only at intermediate levels of confidence. For some participants, we observe a strong divergence at high confidence (Participant 1,4 and 5). This can be understood by the fact that, in this model, 'firing rate' variables can take negative values (in fact, for the steady state in the absence of any input, the firing rate variables take negative values). This leads to extreme value of confidence for long trials. However, the trend in accuracy is not correct for some participants (Participants 1 et 4). Accuracy is an increasing function of confidence (except for participant 5), but the experimental data do not fall within the bootstrapped confidence interval of the simulations, as can be seen in Fig. S2.

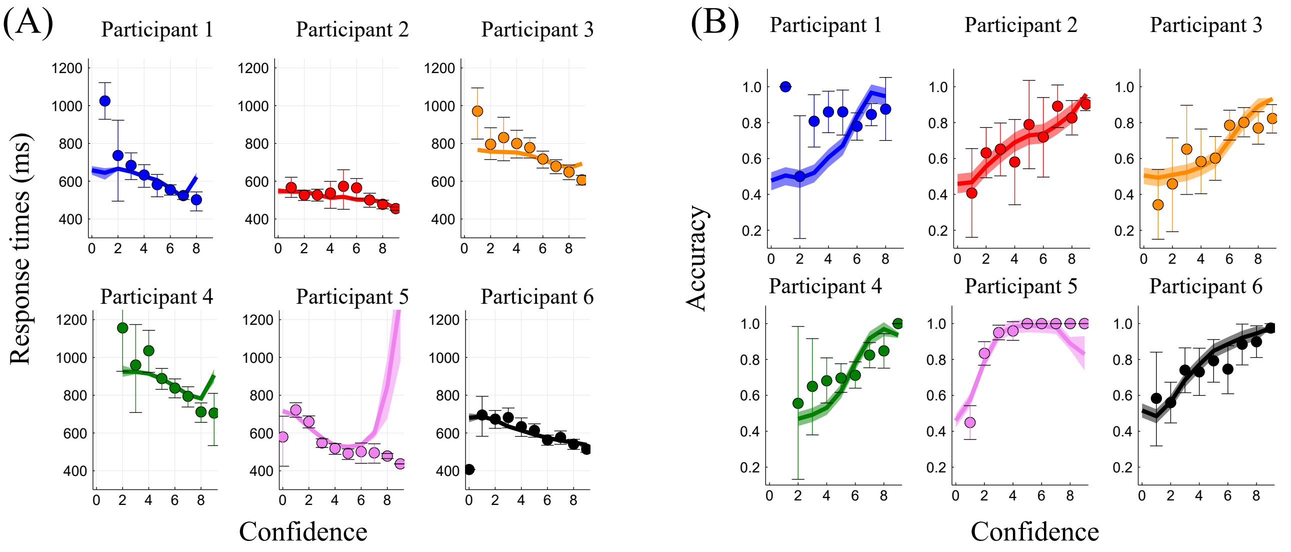


Figure S1: **Response times and Accuracy as a function of confidence in the Usher-McClelland model.** (A) Response times, and (B), Accuracy, with respect to confidence. For both panels: Each sub-panel represents a different participant. The x-axis gives the reported confidence. The dots stand for the experimental data, and the error bar the 95% bootstrapped confidence interval. The line denotes the results of the simulations with the Usher McClelland model.

### S2 - Confidence in success and error trials

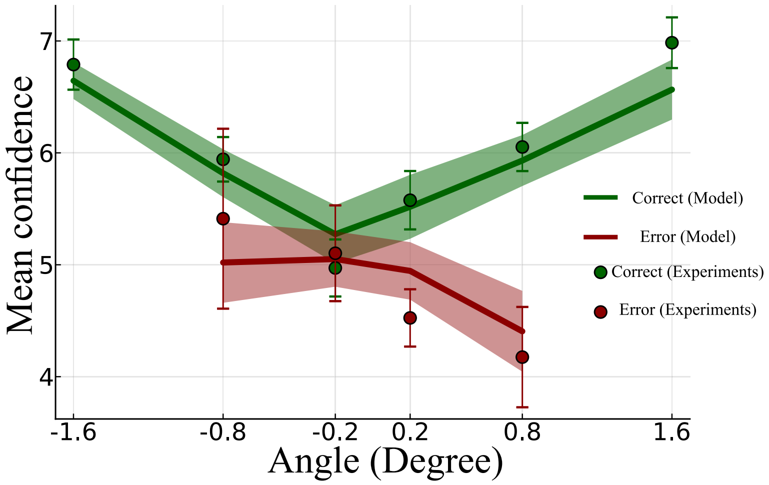


Figure S2: **Confidence as a function of stimulus strength.** We represent the mean confidence as a function of stimulus orientation in correct trials (green), and in error trials (red), for the experimental data (points) and the model simulations (lines). With parameters resulting from the fit on the confidence block, the numerical protocol mimic the experimental one (same number of trials, and same angle values). Due to the discrete levels of confidence, and the high performance in the task, to get enough statistics we combined the data of all subjects. The shaded areas (resp. error bars) denote the $95\%$ bootstrapped confidence interval on the mean for the simulation (resp. data)

### S3 - STATISTICAL ANALYSES

As explained in the paper, we perform a linear mixed effects analysis [Gelman and Hill, 2007] of the history biases of the reaction times, on both the experimental and the numerical simulations data. The linear mixed effects model (LMM) we consider assumes that the logarithm of the response time at trial $n$, $RT_n$, is a linear combination of factors as follows:

$$\ln(RT_n) = a_{0,p} + a_{1,p}|\theta| + a_2 x_{\text{repetition}} + a_{3,p}\ln(RT_{n-1}) + a_4 \text{Conf}_{n-1} \tag{1}$$

with $x_{\text{repetition}}$ a binary variable taking the value 1 if the correct choice for the current trial is a repetition of the previous choice (and 0 otherwise), $\theta$ the orientation of the Gabor (in degree), $RT_{n-1}$ the response times of the previous trials, and $\text{Conf}_{n-1}$ the confidence of the previous trial coded as 0 for *low* and 1 for *high*. The subscript $p$ in a coefficient (e.g $a_{0,p}$) indicates that for this parameter we allow for a random slope per participant.

We present the results obtained with this LMM, equation (1), in Table S2 for the experimental data, and in Table S3 for the numerical data. See the main text, section Results, for the discussion.

We also compare this LMM to other ones that do not include all the terms, using the *ANOVA* function (with the lme4 package [Bates et al., 2015] in RStudio [RStudio Team, 2015]) that performs model comparison based on the Akaike and Bayesian Information Criteria (AIC and BIC) [Bates et al., 2014]. To perform the comparison between the experimental data and the results of the numerical simulations we first transform the response times of each participant using the z-score [Kreyszig, 1979]. This normalization of the response times allows us to study the participants altogether.

We present the results of the models comparison in Table S4. We see that the full model, equation (1), is always to be preferred.

| | Estimate | Std. Error | df | t-value | Pr | |
|---|---|---|---|---|---|---|
| $a_{0,p}$ | 5.428 | 1.466e-01 | 9.0 | 37.038 | $6.92 \cdot 10^{-11}$ | *** |
| $a_{1,p}$ | $-0.1027$ | 0.02390 | 9.0 | $-4.296$ | 0.002001 | ** |
| $a_2$ | $-3.402 \cdot 10^{-2}$ | $2.840 \cdot 10^{-3}$ | $8.472 \cdot 10^3$ | $-11.978$ | $< 2 \cdot 10^{-16}$ | *** |
| $a_{3,p}$ | $1.517 \cdot 10^{-1}$ | $1.651 \cdot 10^{-2}$ | 7.0 | 9.187 | $3.23 \cdot 10^{-5}$ | *** |
| $a_4$ | $-2.063 \cdot 10^{-2}$ | $5.969 \cdot 10^{-3}$ | $5.537 \cdot 10^3$ | $-3.456$ | 0.000553 | *** |

Table S2: **Results of the application of the LMM on the experimental data. We note $**$ for $p < 0.005$ and $***$ for** $p < 0.001$.

| | Estimate | Std. Error | df | t value | Pr | |
|---|---|---|---|---|---|---|
| $a_{0,p}$ | 5.999 | 0.08032 | 4.229 | 74.690 | $9.22 \cdot 10^{-8}$ | *** |
| $a_{1,p}$ | $-0.01744$ | $5.551 \cdot 10^{-4}$ | 2.886 | $-31.420$ | $9.47 \cdot 10^{-5}$ | *** |
| $a_2$ | $-0.1814$ | $8.133 \cdot 10^{-3}$ | $4.822 \cdot 10^3$ | $-22.301$ | $< 2 \cdot 10^{-16}$ | *** |
| $a_{3,p}$ | $-0.02075$ | $1.545 \cdot 10^{-2}$ | 4.628 | $-1.343$ | 0.24139 | |
| $a_4$ | $-0.02324$ | $8.336 \cdot 10^{-3}$ | $4.847 \cdot 10^3$ | $-2.788$ | 0.00533 | ** |

Table S3: **Results of the application of the LMM on the data from the neural network simulations. We note $**$ for** $p < 0.005$ **and $***$ for** $p < 0.001$.

| | Df | AIC | BIC | LogLik. | p value |
|---|---|---|---|---|---|
| $a_{0,p} + a_{1,p}|\theta| + a_2 x_{\text{repetition}} + a_{3,p}\ln(RT_{n-1}) + a_4\text{Conf}_{n-1}$ | 12 | -335 | -254 | 180 | |
| $a_{0,p} + a_{1,p}|\theta| + a_2 x_{\text{repetition}} + a_{3,p}\ln(RT_{n-1})$ | 11 | -324 | -249 | 173 | 0.0003 |
| $a_{0,p} + a_{1,p}|\theta| + a_2 x_{\text{repetition}} + a_4\text{Conf}_{n-1}$ | 8 | -30 | 25 | 23 | <2e-16 |
| $a_{0,p} + a_{1,p}|\theta| + a_2 x_{\text{repetition}}$ | 7 | -4 | -42 | 9 | <2e-16 |
| $a_0 + a_1|\theta| + a_2 x_{\text{repetition}} + a_3\ln(RT_{n-1}) + a_4\ln(\text{Conf}_{n-1})$ | 7 | -225 | -177 | 119 | <2e-16 |
| $a_0$ | 3 | -475 | -495 | -234 | <2e-16 |

Table S4: **LMM tests on Data, models comparison. The first row gives the tests for the LMM used in the main text. The p-values are for the tests based on BIC and AIC Bates et al. [2014] between the LMM used and the one of the corresponding row.**

We test if correlations between confidence and reaction times could explain the fact that the attractor network is not able to reproduce the bias on reaction times due to the reaction time at the previous trial. To do so we consider following LMM:

$$\ln(RT_n) = a_{0,p} + a_{1,p}|\theta| + a_2 x_{\text{repetition}} + a_{3,p}\ln(RT_{n-1}) \quad (2)$$

| | Estimate | Std. Error | df | t value | Pr | |
|---|---|---|---|---|---|---|
| $a_{0,p}$ | 5.937 | 0.0865 | 3.516 | 68.630 | $1.26 \cdot 10^{-6}$ | *** |
| $a_{1,p}$ | $-0.01745$ | $5.530 \cdot 10^{-4}$ | 2.67 | $-31.559$ | $1.64 \cdot 10^{-4}$ | *** |
| $a_2$ | $-0.1821$ | $8.135 \cdot 10^{-3}$ | $4.822 \cdot 10^{3}$ | $-22.388$ | $< 2 \cdot 10^{-16}$ | *** |
| $a_{3,p}$ | $-0.01197$ | $1.678 \cdot 10^{-2}$ | 3.979 | $-0.714$ | 0.5151 | |

Table S5: **Results of the application of the LMM 2 on the data from the neural network simulations. We note** $**$ **for** $p < 0.005$ **and** $***$ **for** $p < 0.001$.

We note that, even when the previous confidence is not taken into account by the LMM, the LMM does not find an effect of previous reaction times. This means that the attractor neural network does not reproduce this effect that is observed in the experimental data.

S4 - PARAMETERS OF THE MODEL

| Block | Parameter | Participant 1 | Participant 2 | Participant 3 | Participant 4 | Participant 5 | Participant 6 |
|---|---|---|---|---|---|---|---|
| Pure Block | $z(Hz)$ | 10.78 | 12.69 | 14.07 | 12.80 | 10.05 | 12.89 |
| | $\Delta z(Hz)$ | (-0.7,+1.75) | (-2.1,+0.175) | (-1.92,+1.75) | (-0.1,+2.275) | (-1.8,2.1) | (-1.05,+2.45) |
| Confidence | $z(Hz)$ | 13.08 | 13.70 | 14.95 | 12.96 | 12.55 | 14.65 |
| Block | $\Delta z$ (Hz) | (-0.18,+2.28) | (-1.93,+0.18) | (-2.45,+1.40) | (-0.17,+2.1) | (-0.35,+2.8) | (-2.1,+0.53) |

Table S6: **Threshold parameter for each participant after fit of the mean accuracy and response times of the *pure* and *confidence* blocks. The ranges $\Delta z$ correspond to one sigma deviation of the likelihood with respect to the corresponding parameter (see the Material and Methods Section).**

| | Type of fit | Pure Block | Confidence Block |
|---|---|---|---|
| Participant 1 | Quadratic | $c_\theta = 554.8 * \theta - 1444 * \theta^2$ | $c_\theta = 650 * \theta 1.3 \times 10^4 * \theta^2$ |
| Participant 2 | Quadratic | $c_\theta = 729.6 * \theta - 9819 * \theta^2$ | $c_\theta = 1056 * \theta - 1.4 \times 10^4 * \theta^2$ |
| Participant 3 | Linear | $c_\theta = 524.4 * \theta$ | $c_\theta = 634.6 * \theta$ |
| Participant 4 | Quadratic | $c_\theta = 269.8 * \theta - 577.6 * \theta^2$ | $c_\theta = 190 * \theta + 2.17 \times 10^4 * \theta^2$ |
| Participant 5 | Quadratic | $c_\theta = 387.6 * \theta + 4188 * \theta^2$ | $c_\theta = 182.4 * \theta + 4.9 \times 10^4 * \theta^2$ |
| Participant 6 | Linear | $c_\theta = 551 * \theta$ | $c_\theta = 1030 * \theta$ |

Table S7: **Calibration of the stimulus strength parameter: best fit for each participant. $\theta$, in radian, stands for the absolute value of the orientation.**

### S5 - Estimation of the non-decision time distribution

We consider that the nondecision time distribution, noted $\rho_{NDT}$, is described by an ex-Gaussian distribution [Verdonck and Tuerlinckx, 2016]:

$$\rho_{NDT}(t) = \frac{\lambda_{NDT}}{2} \exp\left(\frac{\lambda_{NDT}}{2}(2\mu + \lambda_{NDT}\,\sigma^2_{NDT} - 2t)\right) \operatorname{erfc}\left(\frac{\mu + \lambda_{NDT}\,\sigma^2_{NDT} - t}{\sqrt{2}\sigma_{NDT}}\right) \tag{3}$$

with erfc the complementary error function. For such distribution, the mean non decision time, $\langle NDT \rangle$, is given by

$$\langle NDT \rangle = \mu_{NDT} + \frac{1}{\lambda_{NDT}}. \tag{4}$$

As we assume no correlation between response and non decision times, the total (observed) response time distribution, $\rho_{data}$, can be written as the convolution of the decision time distribution, $\rho_{decision}$, with the nondecision time distribution, $\rho_{NDT}$:

$$\rho_{data}(t) = \rho_{decision}(t) * \rho_{NDT}(t) = \int_0^t \rho_{decision}(t-u)\rho_{NDT}(u)\,du \tag{5}$$

(with $*$ standing for the convolution operation). If the decision time distribution is Gaussian, the resulting total distribution is an ex-Gaussian distribution [Grushka, 1972].

Using maximum likelihood estimation, for each subject we fit the empirical response time distribution (all orientations together) by an ex-Gaussian distribution with parameters $\mu_{data}, \lambda_{data}, \sigma_{data}$ (and we thus have $\langle RT \rangle_{data} = \mu_{data} + 1/\lambda_{data}$). For what concerns the model, we find that the decision time distribution of the attractor neuronal network is well fitted by a Gaussian distribution with parameters $\langle RT \rangle_{network}, \sigma_{network}$. We thus model the decision time distribution by the one provided by the attractor network whose parameters have been calibrated as explained above. Hence, we identify the mean and variance of the decision time distribution with the ones of the network: $\mu_{decision} = \langle RT \rangle_{network},\ \sigma_{decision} = \sigma_{network}$.

Taking the characteristic function of both sides of Equation 5, we get:

$$\left(1 - \frac{it}{\lambda_{data}}\right)^{-1} \exp\left(i\mu_{data}t - \frac{1}{2}\sigma^2_{data}t^2\right) =$$
$$\exp\left(i\mu_{decision}t - \frac{1}{2}\sigma^2_{decision}t^2\right)\left(1 - \frac{it}{\lambda_{NDT}}\right)^{-1} \exp\left(i\mu_{NDT}t - \frac{1}{2}\sigma^2_{NDT}t^2\right)$$

We can then identify the terms on both sides of this equations, which, given the use for the decision parameters of the ones of the attractor network, gives the equations

$$\lambda_{NDT} = \lambda_{data}, \tag{6}$$

$$\langle NDT \rangle = \mu_{NDT} + \frac{1}{\lambda_{NDT}} = \langle RT \rangle_{data} - \langle RT \rangle_{network} \tag{7}$$

and

$$\sigma^2_{NDT} = \sigma^2_{data} - \sigma^2_{network}\,. \tag{8}$$

from which we can compute the non-decision time distribution parameters, $\lambda_{NDT}, \mu_{NDT}, \sigma_{NDT}$.

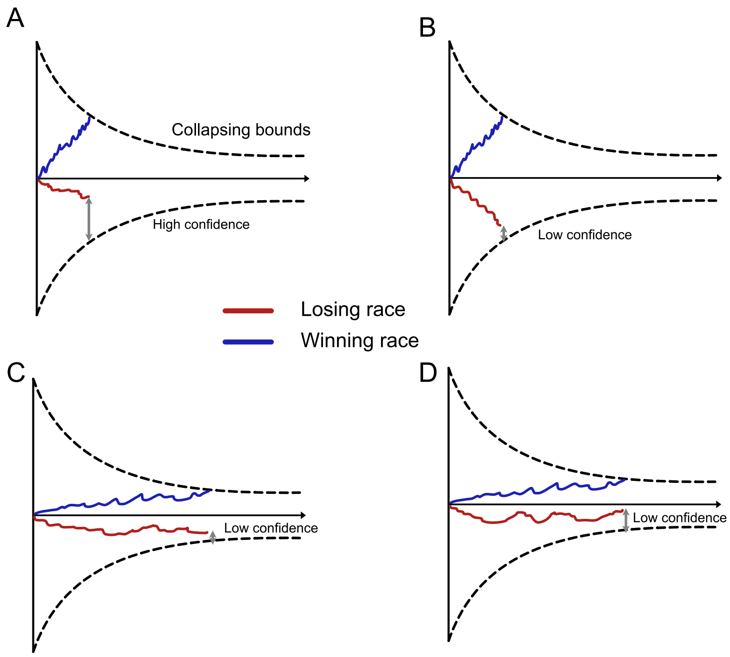


Figure S3: **IRM collapsing bounds.** Each panel represents a different case in the IRM with collapsing bounds. The blue race corresponds to the winning race and the blue one to the losing race. (A) This panel corresponds to the case where the decision was fast and lead to a high confidence trial. (B) Fast decision but low confidence trial. (C) Slow decision and losing race very close to the decision threshold. (D) Slow decision and losing race far to the decision threshold.

#### S6 - DDM WITH COLLAPSING BOUNDS

Figure S3 shows a schematic representation of the IRM with collapsing bounds. The same analysis as the one in the main text for the IRM without collapsing bounds leads to the following conclusions. One can observe two behaviors. Either the decision is fast and it is possible for the model to be in a high-confidence trial, either the decision is slow and the confidence will be low.

If the decision was fast, the model will lead to the opposite sequential effect due to confidence after the relaxation period (similar case as in the main text). However, if the decision was slow the losing race is going to be close to the decision threshold and the confidence will be low for all of these trials. After the relaxation, the losing race will have the same state as in the high-confidence fast trial case. Therefore there will be no sequential effects. This analysis shows that even an IRM with collapsing bounds and a relaxation mechanism will not account for the sequential effects due to confidence that we can model with an attractor neural network.
